# Crystal structures reveal nucleotide-induced conformational changes in G motifs and distal regions in guanylate-binding protein 2

**DOI:** 10.1101/2023.06.28.546747

**Authors:** Sayantan Roy, Bing Wang, Yuan Tian, Qian Yin

**Author notes:** These two authors contribute equally to the manuscript (Sayantan Roy, Bing Wang).

## Abstract

Guanylate-binding proteins (GBPs) are interferon-inducible GTPases that confer protective immunity against a variety of intracellular pathogens including bacteria, viruses, and protozoan parasites. GBP2 is one of the two highly inducible GBPs, yet the precise mechanisms underlying the activation and regulation of GBP2, in particular the nucleotide-induced conformational changes in GBP2, remain poorly understood. In this study, we elucidate the structural dynamics of GBP2 upon nucleotide binding through crystallographic analysis. GBP2 dimerizes upon GTP hydrolysis and returns to monomer state once GTP is hydrolyzed to GDP. By determining the crystal structures of GBP2 G domain (GBP2GD) in complex with GDP and nucleotide-free full-length GBP2, we unveil distinct conformational states adopted by the nucleotide-binding pocket and distal regions of the protein. Our findings demonstrate that the binding of GDP induces a distinct closed conformation both in the G motifs and the distal regions in the G domain. The conformational changes in the G domain are further transmitted to the C-terminal helical domain, leading to large-scale conformational rearrangements. Through comparative analysis, we identify subtle but critical differences in the nucleotide-bound states of GBP2, providing insights into the molecular basis of its dimer-monomer transition and enzymatic activity. Overall, our study expands the understanding of the nucleotide-induced conformational changes in GBP2, shedding light on the structural dynamics governing its functional versatility. These findings pave the way for future investigations aimed at elucidating the precise molecular mechanisms underlying GBP2’s role in the immune response and may facilitate the development of targeted therapeutic strategies against intracellular pathogens.

## Introduction

Cell-autonomous immunity is the collective effector mechanisms that provide protection to both immune and non-immune cells and often induced by proinflammatory cytokines, especially interferons ^1^. Among the thousands of genes induced by interferons is a prominent family of GTPase which accounts for twenty percent of the total population of interferon-stimulated genes ^2, 3^. Interferon-inducible GTPases comprise of four subfamilies: the 65 kDa guanylate-binding proteins (GBPs), the 47 kDa immunity-related GTPases (IRGs), the 72-82 kDa Myxovirus resistant (Mx) proteins, and the 200-285 kDa very large inducible GTPases (VLIGs/GVINs) ^2, 3^.

GBP proteins are widely spread in eukaryotes. A recent hidden Markov modeling revealed that there are 132 intact GBP genes across 32 taxa ^4^. The human GBP family consists of seven members, GBP1-7, all located on chromosome 1. GBPs are composed of a G domain at the N-terminus and a helical region at the C-terminus. Prenylation of certain GBPs at their C-termini CaaX motif brings them to the membrane structures in the cell ^5, 6^. GBPs are most closely related to dynamin family of GTPases that play critical roles in membrane remodeling ^2, 3, 7^. Dynamin family GTPases undergo guanosine nucleotide driven self-assembly to form large homotypic complexes ^8^. Like dynamins, GBPs can homodimerize via their G domains in a parallel fashion ^9, 10^. GBPs possess distinctive features, for example, GBP1 can bind both GTP and GDP with equimolar affinity to produce guanosine-5’-monophosphate (GMP) ^11^. However, the physiological importance of this nucleotide preference is still unknown. Once nucleotide bound, GBP1 exhibit high intrinsic rates of GTPase and GDPase activity that occurs in a structurally conserved two-step reaction ^9^.

Following interferon induction, GBPs confer immunity against a wide range of pathogens. Antibacterial defense was the first function tested across the complete Gbp family ^12^. Subsequent work in the following years established the role of GBPs in restricting apicomplexan parasites ^13^ and certain viruses including human immunodeficiency virus (HIV) and Zika virus ^14^, though the range of antiviral activity is still under characterization. Studies to identify the structures recognized by GBPs to target the pathogen niche have led to the discoveries that GBP5 in mouse macrophages recognizes lipid A moiety of Gram-negative LPS and human GBP1 recognizes LPS O antigen in lung epithelia infected by *Shigella* or *Burkholderia* ^4^. GBP1 can encapsulate several cytosol invasive Gram-negative and Gram-positive pathogens, and has a non-selective nature regarding LPS detection ^12, 15, 16^. Recently, it was shown that GBP2 can also recognize LPS, form LPS aggregates and lead to LPS dependent caspase-4 activation ^16^. Altered self-ligands in the form of liberated intraluminal host ligands could also serve as proxies of infection, apart from conserved microbial structures ^17^.

The myriad of roles played by GBPs in mediating host defense range from forming assembly platforms for caspase-4 recruitment and activation ^18, 19^, promoting the rupture of *Salmonella* containing vacuoles ^20^, disrupting the structural integrity of bacteria ^21^ to pyroptosis in cell ^22^. Out of the three members which are prenylated in the cell, GBP1, 2, and 5, GBP1 received the maximum attention in the last decade and is the most extensively studied, followed by GBP5. Available structures include full-length GBP1 in nucleotide-free and GMPPNP-bound forms ^23, 24^, GBP1 G domain (GBP1GD) in GMPPNP-, GDP·AlFx^−^, GMP·AlFx^−^, and GMP-bound forms ^9^, full-length nucleotide-free GBP1 with farnesyl modification ^25^, a truncated nucleotide-free GBP1 in complex of *Shigella* flexneri effector IpaH 9.8 ^25^, a full-length GDP-bound GBP1 in complex with IpaH 9.8 ^26^, nucleotide-free and GDP·AlFx^−^ bound forms of truncated GBP5, and GDP·AlFx^−^ bound form of GBP5GD ^10^. Recently, the other members of the GBP family, including GBP2, 3, and 4, started gaining importance, when it was shown that GBPs form supramolecular complexes around bacterial surface ^18, 19^. GBP2 is one of the two most highly induced GBPs ^27^ yet it is not well studied as GBP1 or GBP5. Though human GBP1 and GBP2 share sequence identity of 76.3% and similarity of 91.1%, they differ in several noticeable aspects. GBP1 hydrolyzes GTP to GDP and then GMP in two quick successive steps ^9^. While GBP2 is able to produce GMP as the end product, the efficiency is much less. Over 75% of GBP2 hydrolysis products is GDP ^28, 29^. GBP1 binds to GTP, GDP, and GMP in comparable affinities, but GBP2 binding to GMP is ten times lower than that of GTP or GDP ^28^. Furthermore, GDP is a potent inhibitor of GTP hydrolysis by GBP2 but not GBP1 ^28^. It is also unclear how GTP hydrolysis couples with GBP dimerization or resolves after GDP formation and release. Here, we employed crystallographic and biochemical tools to reveal the structural features that underlie the inhibited form of GBP2 and the reason that GBP2 shifts from dimer to monomer upon completion of GTP hydrolysis.

## Results

### GBP2 forms dimer in solution upon GTP hydrolysis

We expressed full-length human GBP2 (1-591) and GBP2 G domain (GBP2GD, 1-309) (Figure 1A) in *E. coli* and purified the proteins to homogeneity. Judging from size-exclusion chromatography, nucleotide-free full-length GBP2 eluted exclusively as a monomer (Figure 1B). Similarly, nucleotide-free GBP2GD eluted mostly as a monomer, but a noticeable population eluted at the position consistent with dimer formation (Figure 1B). Since GBP1 and GBP5 had been shown to form dimers in solution, we proceeded to examine whether and how GBP2 changes oligomeric status when bound to different guanine nucleotides. Binding to the nonhydrolyzable GTP analog GMPPNP or GDP retained GBP2 in monomeric form; however, binding to GDP·AlFx, the GTP hydrolysis transition state mimic, shifted full-length GBP2 completely to the dimer form (Figure 1C). Dimerization depends on the nucleotide binding capability and GTPase activity of GBP2, as an active site mutant, K51A, remained monomeric when incubated with GDP·AlFx (Figure 1C). We then examined whether GBP2GD followed the same pattern of nucleotide binding and dimerization. Interestingly, GMPPNP binding induced full dimer formation of GBP2GD, yet more than half of GBP2GD remained monomeric even in the presence of excessive amount of GDP·AlFx. (Figure 1D). The difference in dimerization patterns suggests that in full-length GBP2 the C-terminal helical domain either contributes additional dimerization interface or allosterically stabilizes GTP hydrolysis-induced GBP2 dimerization at the transition state.

**Figure 1.**
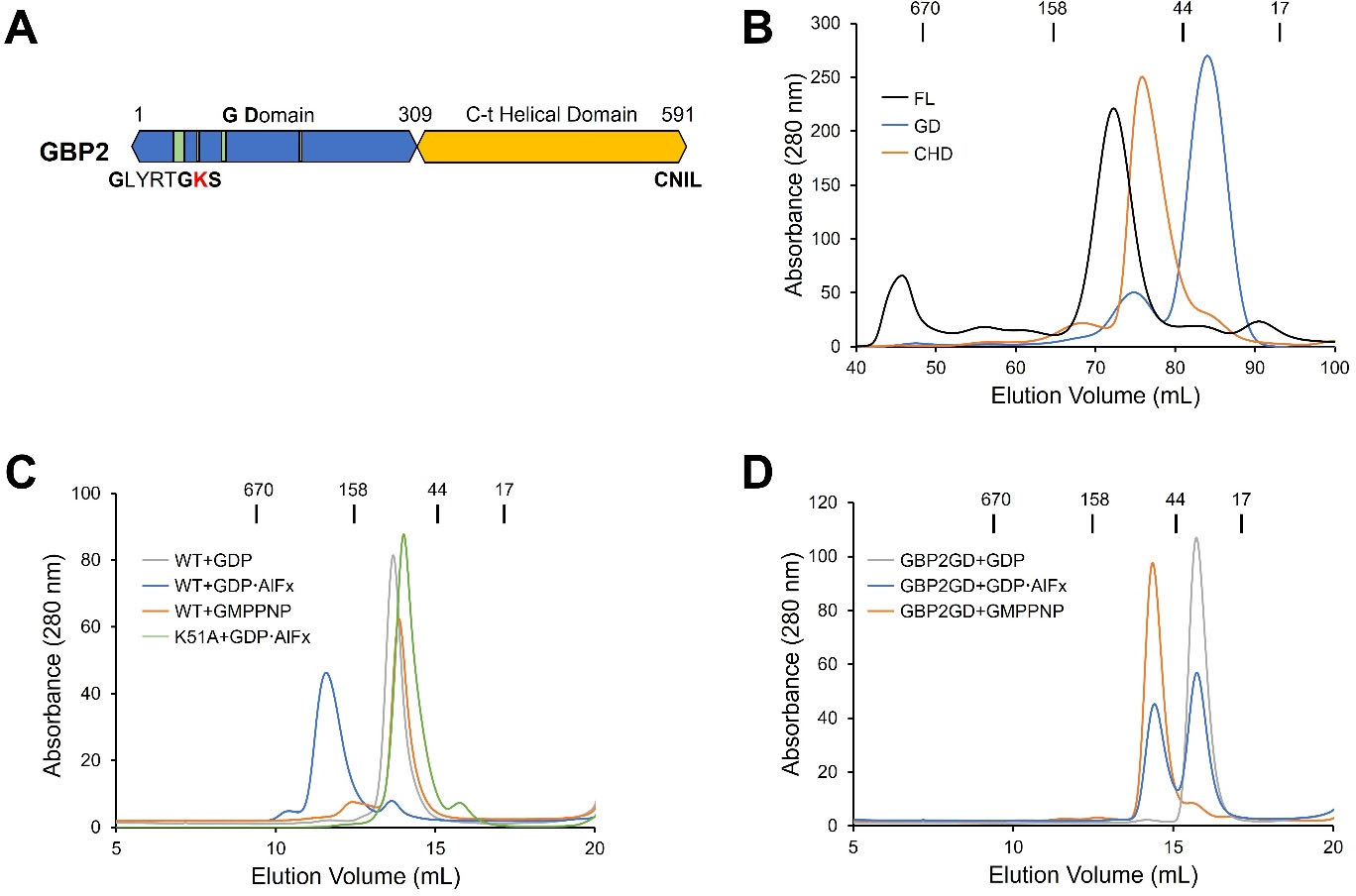
GBP2 forms dimer upon GTP hydrolysis in solution. (**A**) Domain organization of GBP2. G domain is colored blue and C-terminal helical domain is colored orange. G motifs are represented by green vertical bars. G1 motif sequence is spelt out with active site lysine (K) colored in red. The last four residues CNIL in GBP2 constitute an isoprenylation site. (**B**) Size exclusion chromatography (SEC) profiles of full-length GBP2 (FL, black), GBP2 G domain (GD, blue) and C-terminal helical domain (CHD, orange) from a HiLoad 16/600 Superdex200 column. (**C**) SEC profiles of wild-type (WT) GBP2 bound to GDP (gray), GDPꞏAlFx (blue), GMPPNP (orange), and GBP2 K51A mutant incubated with GDPꞏAlFx (green) from a Superdex200 10/300 column. (**D**) SEC profiles of GBP2GD incubated with GDP (gray), GDPꞏAlFx (blue), and GMPPNP (orange) from a Superdex200 10/300 column. In (**B**)(**C**)(**D**) elution positions of protein standards with known MW are marked at the top.

### Crystal structure of GBP2GD in complex with GDP reveals a closed active site

To gain molecular insight into GBP2 activity and dimerization, we determined a 2.1 Å crystal structure of GBP2GD in complex of GDP, which represents the post-hydrolysis, product bound state of GBP2 (Suppl Table 1). The overall structure of GBP2GD contains a mixture of seven beta sheets and seven alpha helices typical of the small GTPases with insertions belonging to the GBP family (Figure 2A and Suppl Figure 1). The guanine base and ribose are enclosed in the space formed by the G1 motif, the G4/RD motif, and the guanine cap (aa 236-255) (Figure 2B and Suppl Figure 1). The G4/RD motif interacts with the guanine moiety via hydrogen bonding. The O6 and N7 groups on the guanine base form dual hydrogen bonds with R181, and N1 and 2’-NH2 are both hydrogen-bonded to D182. P239 adds hydrophobic interaction to the pyrimidine side of the guanine rings. The 2’- and 3’-OH groups of the ribose are held in position by carbonyl oxygens of L245 and A246 in the guanine cap (Figure 2C). At the other end of GDP, the diphosphate group fits snugly into the pocket formed by G1/P-loop, G2/Switch I, and G3/Switch II motifs. Aside from hydrogen bonding with mainchain amines of R48, T49, G50, K51, and S52, the β-phosphate is further elaborately coordinated by side chains of K51, S52, and interestingly, E99. The interactions between the diphosphate group and GBP2 are further reinforced by the hydrogen bonds between the mainchain amines of Y53 and G68 and the α-phosphate (Figure 2D). The nucleotide-binding pocket wraps so tightly around the diphosphate moiety that there is no space to accommodate a magnesium ion even when abundant magnesium was included during crystallization.

**Figure 2.**
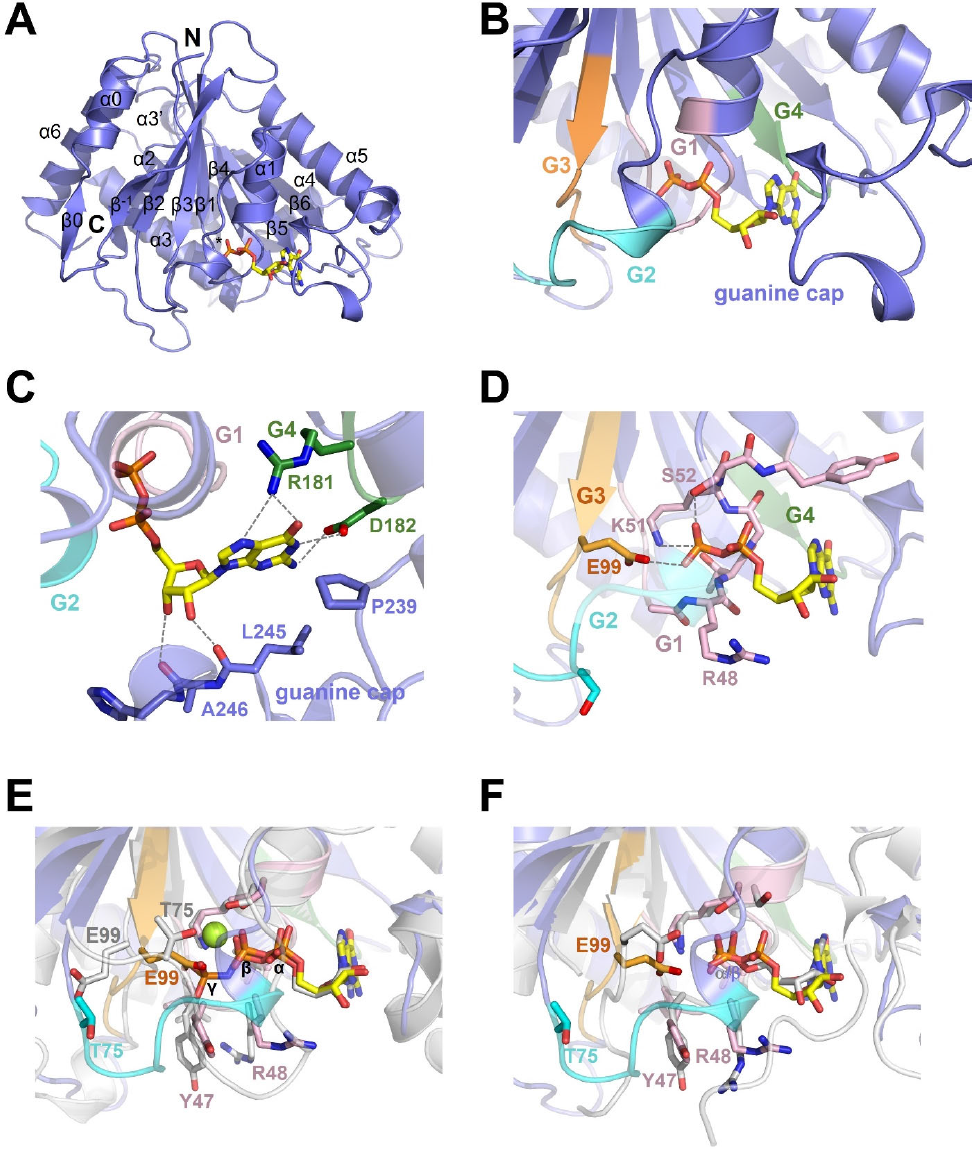
GBP2GD adopts a closed active site conformation when bound to GDP. (**A**) Cartoon representation of GBP2GD·GDP structure with GDP shown as sticks. N- and C-termini of GD and the secondary structure elements are labeled. * indicates helix α4’ at the back of the structure. (**B**) Close-up view of GBP2GD active site when bound to GDP. G1, G2, G3, and G4 motifs are colored in pink, cyan, orange, and green, respectively. (**C**) Close-up view of guanine-ribose-binding space. Critical residues in G4 motif and guanine cap are shown as sticks and hydrogen bonds are represented by dashed gray lines. For clarity, cartoon representation of the rest part of the protein are partially transparent. (**D**) Close-up view of diphosphate-binding pocket. Crucial residues in G1, G2, and G3 motifs are shown as sticks, and hydrogen bonds with side chains are represented by gray dashed lines. G2 motif laying in front of β-phosphate is shown as transparent for clarity purpose. (**E**) Close-up view of superposed nucleotide binding pockets in GBP2GD·GDP structure and GBP1·GMPPNP structure (PDB ID: 1F5N). The residues that display large scale conformational change are shown as sticks with GBP2 residues colored as in (**B**) and GBP1 residues in gray. The magnesium ion in GBP1 structure is represented by a green sphere. (**F**) Close-up view of superposed nucleotide binding pockets in GBP2GD·GDP structure and GBP1GD·GMP structure (PDB ID: 2D4H).

GBP2GD·GDP structure superposes well with that of homologous GBP1 in complex with GMPPNP (GBP1·GMPPNP, PDB ID 1F5N), and the GDP molecule occupies exactly the same space as the GDP portion in GMPPNP (Figure 2E). Nonetheless, we noticed substantial conformational changes in the G2/Switch I region and a distal region on the opposite site of GD (Figure 2E and Suppl Figure 2A), highlighting the two distinct states of GBP2 before and after GTP hydrolysis. In the GBP2GD·GDP structure, G2/Switch I region swings out prominently, with some Cαs moving more than 10 Å away from the bound nucleotide (Suppl Figure 2A). On the opposite side, the most prominent difference lies in α3-α3’ region. While α3 remains in the same position but moves about 3 Å as a whole, α3’ swings ∼ 40° from its equivalent position in GBP1·GMPPNP structures. The N-terminal half of α4’ also rotates ∼ 20°, resulting in 5-7 Å movement of corresponding residues. The last α helix in GD, α6, also shows slight movement at its C-terminus (Suppl Figure 2A).

G1 and G3 motifs remain relatively immobile at first look, but closer examination revealed large scale changes in side-chain orientations that result in the substantial difference in phosphate binding pocket. In the G1 motif, the side chains of Y47 and R48 swings about 70° from their outward orientations in GBP1·GMPPNP structure, pushing away the G2 motif to its current position in the GBP2GD·GDP structure (Figure 2E). E99 in the G3 motif flips 120° inward, occupying the γ-phosphate position in the GMPPNP-bound structure (Figures 2D and 2E). The negative charge of E99 side chain is only 2.6 Å away from the β-phosphate in GDP. Movement of the G2/Switch I motif disrupts the conformation that is essential for magnesium coordination. In GBP1·GMPPNP structure, the side chains of S52 in G1 motif and T75 in G2 motif hold the magnesium ion in place to coordinate with β- and γ-phosphates. In GBP2GD·GDP, T75 moves 7 Å away due to the prominent movement of G2 motif, no longer capable to coordinate magnesium ion together with S52 (Figure 2E). Together with the swinging in of E99, the GDP-binding pocket can no longer accommodate a magnesium ion around diphosphate.

While GBP1 quickly hydrolyzes GTP to the final product of GMP, the major product of GBP2-mediated hydrolysis is GDP ^28, 29^. We then compared GBP2GD·GDP and GBP1GD·GMP structures to examine whether the product-bound structures of GBP1 and GBP2 share more similarities. Though the residues Y47, R48, and E99 display very similar orientations in the two structures (Suppl Figure 2B), G2 motif and guanine cap display substantial movement. In GBP2GD·GDP structure, the active site is more closed. G2 and the region immediately preceding it are ordered and closely packed into the diphosphate moiety. In GBP1GD·GMP structure, the same region swings outwards and towards ribose moiety, not making contacts with the α phosphate, which occupy almost the same position as the β-phosphate in GBP2GD·GDP structure. Part of the G2 motif even becomes disordered and is not visible in the GBP1GD·GMP structure. At the opposite side, α3, α3’, and α4’ all show movement in the two structures, recapitulating the differences we observed between GBP2GD·GDP and GBP1·GMPPNP structures (Suppl Figure 2A).

GBP2 GD is monomeric when bound to GDP and partially dimeric when bound to GDP·AlFx (Figure 1D). We then set out to identify the structural features that account for the monomeric nature of GBP2GD·GDP. Superposing the GBP2GD·GDP structure onto the GBP1GD dimer structure (GBP1GD·GDP·AlFx, PDB ID 2B92) reveals conformations that are incompatible with dimer formation of GBP2GD in the GDP-binding state (Suppl Figure 3A). In the GDP-bound form, the region immediately following switch II (the region connecting β3 and α2) will clash with the N-terminus of α3 of the other protomer (Suppl Figure 3B). In addition, the G2/switch I region moves away from the dimer interface (Suppl Figure 3C), losing contacts with the loop between β5 and α4 of the other protomer. Together, conformational rearrangements in switch I and switch II regions dictate that once GTP is hydrolyzed to GDP, GBP2 can no longer form dimer.

### Mutation of G motif residues abrogated GTPase activity of GBP2

In the GBP2GD·GDP crystal structure, several conserved G1 and G3 residues interact with the diphosphate moiety, including R48, K51, and E99 (Figures 2D). To validate our observation in the crystal, we mutated the residues in both GBP2FL and GBP2GD and evaluated the activity of the mutants in comparison to wild-type proteins using the colorimetric malachite green assay. Individual mutation of G1 motif residues R48 or K51 to alanine (R48A and K51A) completely abrogated the GTPase activity of both GBP2FL and GBP2GD (Figures 3A and 3B). As a control, mutation of a G2 residue S73 that points away from GDP in the structure (S73A) did not reduce the GTPase activity in GBP2 (Figures 3A and 3B). We are particularly interested in the G3 motif residue E99 since its negative charge is positioned right next to the β-phosphate in the crystal structure (Figure 2D). Mutation of E99 to glutamine (E99Q) reduced GTPase activity to ∼ 40% of the wild type but did not completely abolish it (Figures 3A and 3B), suggesting the negative charges on E99 are optimal for GTPase activity, but hydrogen bonding still function at a lower level. Mutation of E99 to leucine or aspartic acid (E99L and E99D) both abrogated GBP2 activity (Figures 3A and 3B), suggesting the negative charges of E99 must be precisely positioned in the active site for GTPase activity.

**Figure 3.**
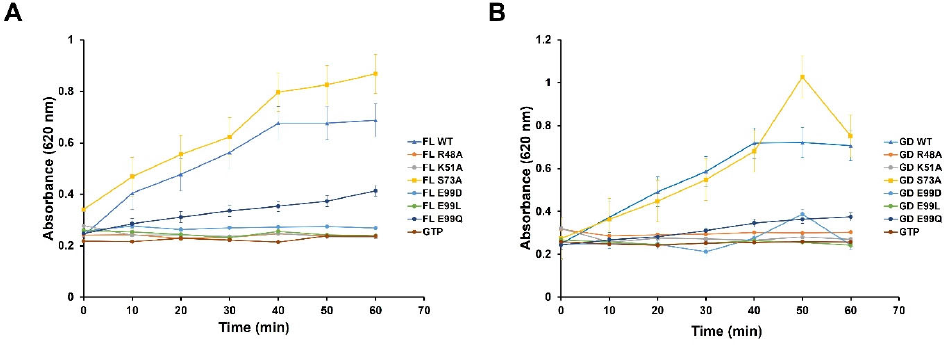
Mutation of critical G motif residues abrogated GTPase activity of GBP2. (**A**) Comparison of the GTPase activities of wild-type (WT) and mutant full-length GBP2. (**B**) Comparison of GTPase activities of wild-type (WT) and mutant GBP2GD. The graphs shown in (**A**) and (**B**) are representatives of at least two independent experiments. The value plotted is an average of duplicate measurements in the same experiment.

When conducting GTPase assays, we noticed that at the same concentration, GBP2GD displayed slightly higher activity than GBP2FL (Suppl Figure 4A), a phenomenon also observed for GBP1 ^9^. We then investigated whether the CHD domain affects GD activity *in trans*. GBP2GD activity remained the same in the presence of increasing amount of GBP2CHD (Suppl Figure 4B), suggesting isolated CHD does not substantially regulate GD enzymatic activity.

### Crystal structure of nucleotide-free full-length GBP2 reveals structural plasticity of CHD

GBP2FL and GBP2GD manifested different dimerization patterns upon nucleotide binding but no drastic differences in GTP hydrolysis (Figures 1C, 1D, and Suppl Figure 4A). We then structurally characterized GBP2FL to uncover any structural basis that accounts for such difference, in particular the effect of the C-terminal helical domain on the G domain. The expression level of full-length GBP2 was low, making structural studies difficult. However, mutation of the active site lysine to alanine (K51A) substantially increased GBP2 yield. We successfully crystallized the full-length GBP2 K51A mutant (GBP2^K51A^) and determined and refined the structure to 2.7 Å (Suppl Table 1). Though abundant GDP and magnesium ions were present during crystallization, no GDP or magnesium ion density was observed in the structure. The structure thus likely represents the nucleotide-free form of full-length GBP2. The GBP2^K51A^ structure assumes an elongated shape, with the globular GTPase domain at one end and the C-terminal helical domain extending out for about 90 Å (Figure 4A). The C-terminal helical domain is composed of six α helices α7 – α13 (Figure 4A and Suppl Figure 1). α7, α8, and the N-terminal half of α9 loosely form one three-helix bundle, and the C-terminal half of α9, α10, and α11 form another three-helix bundle. The region consisting of α7 to α11, also known as the middle domain (MD) ^23^, does not contact the G domain at all. The second-to-last alpha helix α12 folds back towards the GTPase domain, and the last helix α13 makes another about turn to reach towards the center of the helical region. Together, the C-terminal ∼ 15 residues of α12 and α13 make contacts with α3’ and α4’ on the GTPase domain, which localize on the opposite side of the nucleotide binding site. The GD-CHD interface is mostly hydrophilic. The most prominent electrostatic interaction is formed between E554 and E561 on α12 and R225 on α4’, E561 and K205 at the end of α4, and E554 and R231 on β6 (Figures 4B and 4C). In addition, E543 on α12 forms hydrogen bonds with the mainchain NH groups of G282 and G283 between the last two helices (α5 and α6) of the G domain, and the sidechain NH group of K551 on α12 forms a hydrogen bond with the mainchain carbonyl oxygen of A167 on α3’ (Suppl Figure 5A). A second patch of interactions is at the junction of GD and CHD, between α6 on the GD and the N-terminus of α7. In particular, E321 on α7 forms salt bridges with R290 on α6 (Suppl Figure 5B).

**Figure 4.**
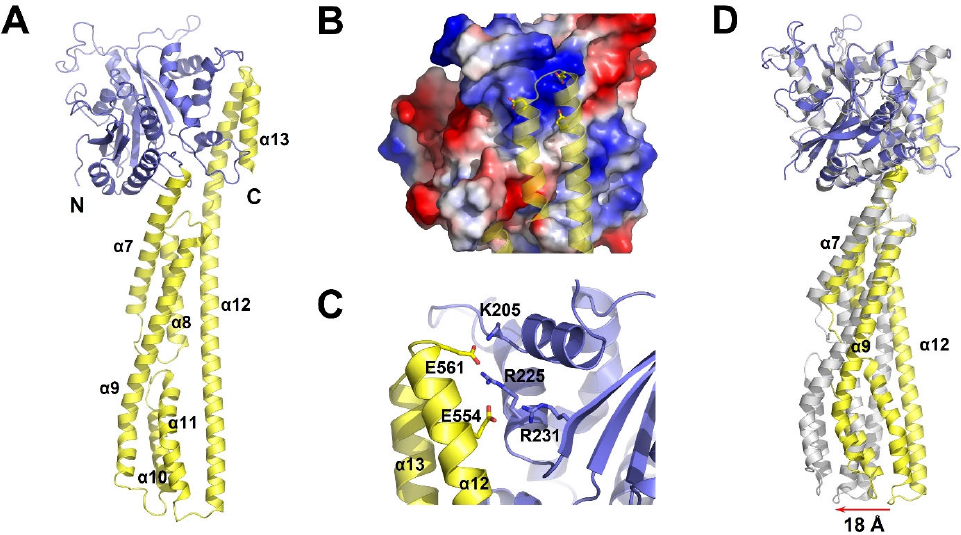
Full-length GBP2 structure displays flexibility between the G domain (GD) and the C-terminal helical domain (CHD) (**A**) Cartoon representation of full-length GBP2^K51A^ structure. The G domain is colored blue as in Figure 2. The C-terminal helical domain is colored yellow. Secondary structure elements in CHD are labeled. (**B**) The interactions between GD and CHD are mainly electrostatic. α12-α13 helices are shown as cartoon, while the GD is shown as the electrostatic surface. (**C**) A close-up view of the GD-CHD interface showing the major electrostatic interactions. Key residues are labeled. (**D**) Superposition of nucleotide-free GBP2^K51A^ structure (blue and yellow) with nucleotide-free GBP1 structure (PDB ID: 1DG3, gray). Orientation of the molecules are rotated by 40 degrees compared to (**A**) to highlight the movement. The red arrow marks the movement of the far end of the CHD.

Overall, the GBP2^K51A^ structure highly resembles the structure of wild-type GBP2 in the nucleotide-free form (PDB ID 7E58) ^10^. Superposition of the two structures reveals an RMSD of 0.772 Å over 477 Cα (out of 545 common Cαs) (Suppl Figure 6A). The two GD domains superpose with an RMSD of 0.488 Å over 243 Cα (out of 276 common Cαs) ^10^. The major deviation comes from the active site (Suppl Figure 7A). While the G1 motif displays modest structural variations, the G2/switch I region, G3/switch II region, and the guanine cap all show large scale movement (Suppl Figure 7A). Similarly, the GD domain in the GBP2^K51A^ structure is highly similar to the GD structures in the two available full-length GBP1 structures, with RMSD values of 0.543 Å over 232 Cα (out of 268 common Cαs with nucleotide-free GBP1, PDB ID: 1DG3) and 0.628 Å over 238 Cα (out of 296 common Cαs, GMPPNP-bound GBP1, PDB ID: 1F5N). Structural deviations in the GD domains cluster in the G motifs and the guanine cap (Suppl Figures 7B-C), consistent with a flexible active site in the absence of substrates or products. The C-terminal helical domain, however, displays remarkable swinging movement. Compared to the nucleotide-free GBP1 structure, the distal end of the C-terminal helical region moves a striking 18 Å (Figure 4D). While the N-terminal half of α7, the C-terminal third of α12, and the whole α13 superpose well with their counterparts in GBP1, tilting of the C-terminal helical region is collectively mediated by the varied bending in the middle of α7, tilting angle of the N-terminus of α9, and the bending in α12 around T530 (Figure 4D). Indeed, the large-scale swinging movement of the C-terminal helical domain has been observed for full-length GBPs and GBP proteins lacking the last two α helices (Suppl Figure 6) ^23–25^. Superposing available GBP2 structures reveals conformational changes in the α3-α3’ linker and α4’ region (Suppl Figure 8), which likely contribute to the movement of the C-terminal helical domain.

## Discussion

Using recombinant proteins and guanine nucleotide mimics, we showed that GBP2FL is monomeric in both nucleotide-free and substrate-binding forms. It dimerizes at the transition state of GTP hydrolysis, then returns to the monomeric form when only GDP, the product, remains binding (Figure 1C). GBP2GD, however, displays a different profile. GBP2GD is readily dimeric when bound to the substrate mimic GMPPNP but shows a ∼ 50:50 split between dimers and monomers at the transition state (Figure 1D). As GBP2GD and GBP2FL displayed comparable GTPase activity but contrasted in the dimer formation upon GTP hydrolysis (Suppl Figure 4A, Figure 1C-D), we reason that the C-terminal helical domain does not allosterically regulate GD activity. Instead, the C-terminal helical domain further stabilizes the dimer mediated by GD, likely through a secondary dimerization interface. Indeed, a truncated dimeric GBP5 structure shows additional dimerization interfaces in the CHD when bound to GDP·AlFx^−^ ^10^.

We did not observe any magnesium ion in the GBP2GD·GDP structure even though 5 mM magnesium ions were present during crystallization. As magnesium is essential for GBP2 activity (Suppl Figure 4C), we propose that the captured crystal structure represents the second to last step in GTP hydrolysis – magnesium ion already released, but GDP remains binding (Figures 5C, 5D). The presence of GDP in the active site likely induces a low-energy conformation that precludes binding of magnesium ions. Such binding modes may explain the inability of GBP2 to hydrolyze GDP, even when GBP2 is capable to directly hydrolyze a small portion of GTP to GMP^28^. Any released GDP intermediate will function more as an inhibitor than a substrate. GBP1 cannot utilize GDP as a substrate either, but GDP does not inhibit the hydrolysis of GTP to GMP, likely because GTP binds to GBP1 with higher affinity and is quickly hydrolyzed to GMP without releasing the GDP intermediate or the magnesium ion. In the crystal structure, the negatively charged side chain of E99 points towards the β phosphate of GDP, also negatively charged, seemingly creating a charge incompatibility. We argue that this repulsion between the negative charges facilitates the release of GDP and prepares GBP2 for the next round of GTP hydrolysis. Indeed, mutating E99 to the neutral residue Q reduced GBP2 activity by 60% (Figure 3).

**Figure 5.**
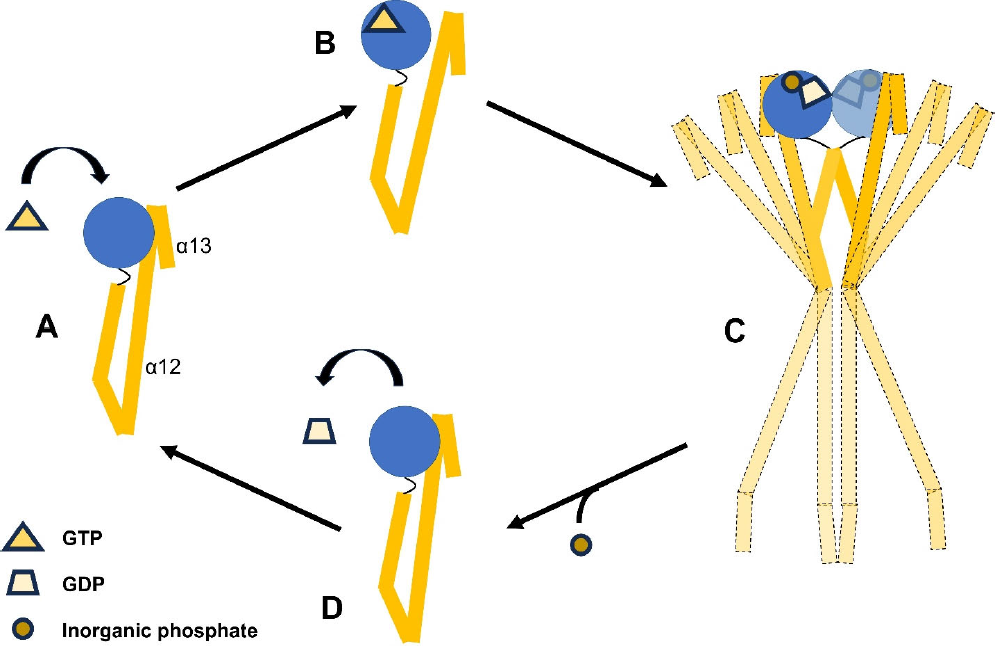
Diagram of GBP2 conformational changes through a complete GTP hydrolysis cycle in solution. GTP binding to the G domain (GD, blue sphere) (**A**) prepares the dissociation of α12 from GD (**B**). (**C**) GTP hydrolysis leads to GBP “cross-over” dimerization at the transition state. α12-α13 may assume orientations ranging from contact with the GD in the other protomer to extending out (represented by dashed outlines). Release of the inorganic phosphate returns GBP2 to monomeric form (**D**). After GDP dissociation, GBP2 returns to the nucleotide-free state (**A**) for next round of GTP binding and hydrolysis.

GBP2 belongs to the family of p67 GTPases with all the signature GTP binding motifs (Suppl Figure 1). Residue K51 resides in the G1 motif, also known as the P-loop (Suppl Figure 1), which contributes to nucleotide binding and positioning ^30, 31^. Mutation of lysine 51 to alanine (K51A) neutralized the GTP hydrolysis enzymatic activity and disrupted the nucleotide dependent oligomerization of GBP2 (Figure 1C, Figure 3A and 3B). R48, conserved in all GBP members, serves the purpose of the catalytic arginine finger, which usually is provided *in trans* by the GTPase activating proteins (GAPs) as in the case of Ras ^32^. R48A mutation abolished the enzymatic activity of GBP2, which is in accordance with what has been observed for GBP1 previously ^33^. The presence of R48 explains the intrinsically high GTPase activity of GBP2 in the absence of GAPs. Mutants in full-length GBP2 and GBP2GD showed similar reduction in activity, suggesting the elongated C-terminal helical domain does not have any additional contribution to nucleotide binding or GTP hydrolysis.

Overall, our findings suggest a working model that GBP2 undergoes large scale conformational changes during GTP hydrolysis cycle (Figure 5). GTP binding and hydrolysis induces conformational changes in the G domain (GD), both in the nucleotide binding region and the α3, α3’, and α4 region on the opposite side of the nucleotide binding region. Conformational changes in the GD enable GD dimerization. They are further transmitted to the CHD via the α3-α3’ linker, leading to α12 moving away from the GD (Figures 5A-B). Dissociation of α12 from GD substantially extends the range of CHD movement, which favors the formation of a cross-over structure in the transition state ^10^. A previous model suggested a possible conformation of full-length GBP2 dimer in the transition state, where α12 and α13 of CHD of one protomer interacts with the GD of the other protomer ^10^ (Figure 5C). However, no full-length dimeric GBP structure is available to substantiate such a closed dimer conformation. Recent studies showed that farnesylated GBP1 can attain an open conformation in the presence of the transition state analog, where the protein assumes an elongated shape with α12 and α13 stretching out instead of interacting with the GD (Figure 5C). Such elongated structures are necessary for coating bacterial membranes as has been shown for GBP1 ^15^. It is unclear whether farnesylation and interaction with bacterial membrane are essential for the formation of such elongated structures. We propose that in solution, α12 and α13 are capable of assuming any possible conformation in dimeric GBP2, from closely interacting with the GD of the other protomer to fully stretching out (Figure 5C). Once the bound GTP is hydrolyzed, the inorganic phosphate is released, leading to a GDP-bound conformation that dissociates the GD-mediated GBP2 dimer. The final step of GDP release returns GBP2 to the nucleotide-free form, completes one cycle of GTP hydrolysis, and prepares GBP2 for the next round of GTP binding and hydrolysis (Figure 5D).

## Materials and Methods

### Molecular cloning and site-directed mutagenesis

Constructs of full-length human GBP2 (GBP2FL, 1-591), GBP2 G domain (GBP2GD, 1-309) were amplified from the cDNA (Accession number: BC073163) using standard polymerase chain reaction, digested with *BamH*I and *Sal*I, and subcloned into a modified pRSFDuet-1 vector containing an N-terminal His_6_-SUMO tag. All mutants of GBP2FL and GBP2GD were generated by Agilent QuikChange Site-Directed Mutagenesis Kit following the manufacturer’s protocol. The primers used for cloning and mutagenesis are listed in Supplementary Table 2. GBP2GD mutants were generated using the same primers for the respective GBP2FL mutants. All constructs were sequenced (Eurofins) to ensure the correct sequences and open-reading frames.

### Protein expression and purification

GBP2FL was overexpressed in *Escherichia coli* BL21-CodonPlus (DE3)-RIPL cells (Agilent). The cells were grown in LB broth containing 34 µg/ml chloramphenicol and 50 µg/ml kanamycin at 37 °C. At OD_600_ of 0.7, the cells were induced with 0.4 mM isopropyl β-D-1-thiogalactopyranoside (IPTG, GoldBio). Post induction, the cells were grown at 20°C for 16-20 hours. After incubation, the cells were harvested and resuspended in Lysis Buffer (50 mM sodium phosphate pH 7.4, 300 mM NaCl, 20 mM imidazole, 10% (v/v) glycerol, and 5 mM β-mercaptoethanol). The resuspended cell pellet was sonicated and centrifuged at 31000 × *g* at 4 °C for 45 min. The supernatant was loaded onto a gravity column containing Ni-NTA beads (Qiagen) equilibrated with Lysis Buffer. Ni-NTA beads were then washed with 10 column volumes of Wash Buffer (50 mM sodium phosphate pH 7.4, 300 mM NaCl, 30 mM imidazole, and 5 mM β-mercaptoethanol). Bound proteins on the beads were eluted with Elution Buffer (50 mM sodium phosphate pH 6.0, 300 mM NaCl, 10% (v/v) glycerol, and 300 mM imidazole). The eluted protein fractions were pooled and treated with the Ulp1 protease at 4 °C overnight to cleave the His_6_-SUMO tag. The overnight treatment also included 25 mM EDTA to remove any endogenous magnesium ion and guanine nucleotides. Concomitant with Ulp1 cleavage, the eluted proteins were dialyzed against Dialysis Buffer (20 mM Tris pH 8.0, 150 mM NaCl, and 5 mM β-mercaptoethanol) to remove imidazole. The cleaved His_6_-SUMO tag and uncleaved His_6_-SUMO-GBP2 were separated from tag free GBP2 by passing the solution over a His-Trap column (GE Healthcare/Cytiva). Tag free GBP2 in the flow-through was concentrated and subjected to size exclusion chromatography on a HiLoad16/600 Superdex 200pg column (GE Healthcare/Cytiva) using the running buffer (20 mM Tris pH 8.0, 150 mM NaCl, and 5 mM β-mercaptoethanol). Peak fractions corresponding to the protein were pooled, concentrated, aliquoted, flash frozen in liquid nitrogen, and stored at −80 °C. GBP2GD and all mutants were expressed and purified in the same way.

### Analytical size exclusion chromatography

Oligomerization states of GBP2FL, GBP2GD, and GBP2FL^K51A^ were analyzed by size exclusion chromatography using a Superdex 200 increase 10/300 GL column (GE Healthcare/Cytiva). 20 µM protein was incubated at 4 °C overnight with 200 µM of the corresponding nucleotide. For the transition state intermediate, 300 µM AlCl_3_ (Sigma) and 10 mM NaF (Sigma) were added along with 200 µM GDP (guanosine 5’-diphosphate sodium salt, Sigma). The column was equilibrated with the running buffer containing 20 mM Tris-HCl pH 8.0, 150 mM NaCl, 5 mM MgCl_2_ and 2 mM Tris (2-carboxyethyl) phosphine (TCEP, GoldBio).

### Crystallization and Structural Determination

Crystallization screening was carried out using a Crystal Gryphon Crystallization Robot (Art Robbins Instruments) with 96-well intelli 3 well crystallization plate (Hampton Research). 0.3 µl protein solution was mixed with 0.3 µl reservoir solution and was set up as a sitting drop with 100 µl reservoir solution at 16 °C. For crystal optimization, 1 µl of protein was mixed with 1 µl of reservoir solution. Both GBP2GD·GDP and GBP2FL^K51A^ crystals were obtained by hanging drop vapor diffusion method at 16 °C. GBP2GD at 13 mg/ml was incubated overnight with 10-fold molar excess of GDP and 5 mM MgCl_2_ before mixing with the reservoir solution of 0.04 M KH_2_PO4, 12% PEG 8000, and 18% glycerol. GBP2FL^K51A^ at 12.1 mg/ml was mixed with the reservoir solution of 0.7 M NaH_2_PO_4_, 1.05 M K_2_HPO_4_, and 0.1 M sodium acetate pH 3.8. Crystals were flash frozen in respective reservoir solution supplemented with glycerol to the final concentration of 30% (v/v) as the cryoprotectant.

Diffraction data were collected at National Synchrotron Light Source II (NSLS II) beamline AMX 17-ID-1. Diffraction data were indexed, integrated, and scaled using HKL2000 ^34^. Both GBP2GD·GDP and GBP2FL^K51A^ structures were solved by molecular replacement using Phenix ^35^. The search model for GBP2GD·GDP is the structure of GBP1GD·GMP (PDB ID 2D4H), while the search model for GBP2FL^K51A^ is the nucleotide free GBP1 structure (PDB ID 1DG3). Further iterative modeling building and refinement were carried out using Phenix and Coot ^35, 36^.

### GTPase activity assay

For measuring the GTPase activity, malachite green phosphate detection kit (Sigma, Cat No. MAK307) was used. Unless otherwise specified, 500 nM GBP2FL or 300 nM GBP2GD protein (wild type or mutant) was mixed with 200 µM GTP (Sigma, Cat No. G8877) in a reaction buffer containing 20 mM Tris pH 8.0, 150 mM NaCl, 8 mM MgCl_2_ and 1 mM EDTA at room temperature. Aliquots were taken out at indicated time points up to 60 min and the GTP hydrolysis reaction was stopped by 25 mM EDTA. Reagent A and Reagent B from the kit were mixed in 100:1 ratio. 20 μL of the prepared reagent was mixed with 80 μL of EDTA-stopped reaction mixture, added to 96 well clear plate, and incubated at room temperature for 30 min for malachite green color change to develop. Absorbance was read in the plate reader (SpectraMax iD5, Molecular Devices) at 620 nm.

## Acknowledgments

We thank the beamline scientists at NSLSII AMX 17-ID-1 AMX and Dr. T. “Soma” Somasundaram at Institute of Molecular Biophysics, Florida State University for assistance in X-ray crystallographic diffraction data collection. This work was supported by FSU startup funds and National Institutes of Health grants R00AI108793 and R01AI146330 to Q. Y. We thank members of Yin group for critical readings of the manuscript and helpful discussions.

## Author Contributions

Y.T. and Q.Y. conceived the initial experimental plan. S.R. and Y.T. expressed and purified the proteins. S.R., B.W., Y.T., Q.Y. crystallized the proteins and determined the structures. S.R. carried out GTPase activity assays. S.R. and Q.Y. drafted the manuscript and all authors edited the manuscript.

## Data availability

Coordinates and structural factors for human GBP2GDꞏGDP and human GBP2FL^K51A^ structures have been deposited in the Protein Data Bank with the accession codes 6VKJ and 7M1S, respectively.

## References

1 Randow, F., MacMicking, J. D. & James, L. C. Cellular self-defense: how cell-autonomous immunity protects against pathogens. Science 340, 701–706, doi:10.1126/science.1233028 (2013).

2 Kim, B. H., Shenoy, A. R., Kumar, P., Bradfield, C. J. & MacMicking, J. D. IFN-inducible GTPases in host cell defense. Cell host & microbe 12, 432–444, doi:10.1016/j.chom.2012.09.007 (2012).

3 Martens, S. & Howard, J. The interferon-inducible GTPases. Annual review of cell and developmental biology 22, 559–589, doi:10.1146/annurev.cellbio.22.010305.104619 (2006).

4 Tretina, K., Park, E. S., Maminska, A. & MacMicking, J. D. Interferon-induced guanylate-binding proteins: Guardians of host defense in health and disease. The Journal of experimental medicine 216, 482–500, doi:10.1084/jem.20182031 (2019).

5 Stickney, J. T. & Buss, J. E. Murine guanylate-binding protein: incomplete geranylgeranyl isoprenoid modification of an interferon-gamma-inducible guanosine triphosphate-binding protein. Molecular biology of the cell 11, 2191–2200, doi:10.1091/mbc.11.7.2191 (2000).

6 Nantais, D. E., Schwemmle, M., Stickney, J. T., Vestal, D. J. & Buss, J. E. Prenylation of an interferon-gamma-induced GTP-binding protein: the human guanylate binding protein, huGBP1. J Leukoc Biol 60, 423-431, doi:10.1002/jlb.60.3.423 (1996).

7 Daumke, O. & Praefcke, G. J. Invited review: Mechanisms of GTP hydrolysis and conformational transitions in the dynamin superfamily. Biopolymers 105, 580–593, doi:10.1002/bip.22855 (2016).

8 Praefcke, G. J. & McMahon, H. T. The dynamin superfamily: universal membrane tubulation and fission molecules? Nature reviews. Molecular cell biology 5, 133–147, doi:10.1038/nrm1313 (2004).

9 Ghosh, A., Praefcke, G. J., Renault, L., Wittinghofer, A. & Herrmann, C. How guanylate-binding proteins achieve assembly-stimulated processive cleavage of GTP to GMP. Nature 440, 101–104, doi:10.1038/nature04510 (2006).

10 Cui, W. et al. Structural basis for GTP-induced dimerization and antiviral function of guanylate-binding proteins. Proceedings of the National Academy of Sciences of the United States of America 118, doi:10.1073/pnas.2022269118 (2021).

11 Praefcke, G. J., Geyer, M., Schwemmle, M., Robert Kalbitzer, H. & Herrmann, C. Nucleotide-binding characteristics of human guanylate-binding protein 1 (hGBP1) and identification of the third GTP-binding motif. Journal of molecular biology 292, 321–332, doi:10.1006/jmbi.1999.3062 (1999).

12 Kim, B. H. et al. A family of IFN-gamma-inducible 65-kD GTPases protects against bacterial infection. Science 332, 717–721, doi:10.1126/science.1201711 (2011).

13 Fisch, D. et al. Human GBP1 Differentially Targets Salmonella and Toxoplasma to License Recognition of Microbial Ligands and Caspase-Mediated Death. Cell reports 32, 108008, doi:10.1016/j.celrep.2020.108008 (2020).

14 Braun, E. et al. Guanylate-Binding Proteins 2 and 5 Exert Broad Antiviral Activity by Inhibiting Furin-Mediated Processing of Viral Envelope Proteins. Cell reports 27, 2092–2104 e2010, doi:10.1016/j.celrep.2019.04.063 (2019).

15 Zhu, S. et al. Cryo-ET of a human GBP coatomer governing cell-autonomous innate immunity to infection. bioRxiv, 2021.2008. 2026.457804 (2021).

16 Dickinson, M. S. et al. LPS-aggregating proteins GBP1 and GBP2 are each sufficient to enhance caspase-4 activation both in cellulo and in vitro. Proceedings of the National Academy of Sciences of the United States of America 120, e2216028120, doi:10.1073/pnas.2216028120 (2023).

17 Huang, S., Meng, Q., Maminska, A. & MacMicking, J. D. Cell-autonomous immunity by IFN-induced GBPs in animals and plants. Current opinion in immunology 60, 71–80, doi:10.1016/j.coi.2019.04.017 (2019).

18 Wandel, M. P. et al. Guanylate-binding proteins convert cytosolic bacteria into caspase-4 signaling platforms. Nature immunology 21, 880-+, doi:10.1038/s41590-020-0697-2 (2020).

19 Santos, J. C. et al. Human GBP1 binds LPS to initiate assembly of a caspase-4 activating platform on cytosolic bacteria. Nat Commun 11, 3276, doi:10.1038/s41467-020-16889-z (2020).

20 Meunier, E. et al. Caspase-11 activation requires lysis of pathogen-containing vacuoles by IFN-induced GTPases. Nature 509, 366–370, doi:10.1038/nature13157 (2014).

21 Man, S. M. et al. IRGB10 Liberates Bacterial Ligands for Sensing by the AIM2 and Caspase-11-NLRP3 Inflammasomes. Cell 167, 382–396 e317, doi:10.1016/j.cell.2016.09.012 (2016).

22 Pilla, D. M. et al. Guanylate binding proteins promote caspase-11-dependent pyroptosis in response to cytoplasmic LPS. Proceedings of the National Academy of Sciences of the United States of America 111, 6046–6051, doi:10.1073/pnas.1321700111 (2014).

23 Prakash, B., Praefcke, G. J., Renault, L., Wittinghofer, A. & Herrmann, C. Structure of human guanylate-binding protein 1 representing a unique class of GTP-binding proteins. Nature 403, 567–571, doi:10.1038/35000617 (2000).

24 Prakash, B., Renault, L., Praefcke, G. J., Herrmann, C. & Wittinghofer, A. Triphosphate structure of guanylate-binding protein 1 and implications for nucleotide binding and GTPase mechanism. The EMBO journal 19, 4555–4564, doi:10.1093/emboj/19.17.4555 (2000).

25 Ji, C. et al. Structural mechanism for guanylate-binding proteins (GBPs) targeting by the Shigella E3 ligase IpaH9.8. PLoS pathogens 15, e1007876, doi:10.1371/journal.ppat.1007876 (2019).

26 Ye, Y., Xiong, Y. & Huang, H. Substrate-binding destabilizes the hydrophobic cluster to relieve the autoinhibition of bacterial ubiquitin ligase IpaH9.8. Commun Biol 3, 752, doi:10.1038/s42003-020-01492-1 (2020).

27 Boehm, U. et al. Two families of GTPases dominate the complex cellular response to IFN-gamma. J Immunol 161, 6715–6723 (1998).

28 Neun, R., Richter, M. F., Staeheli, P. & Schwemmle, M. GTPase properties of the interferon-induced human guanylate-binding protein 2. FEBS letters 390, 69–72 (1996).

29 Rajan, S., Pandita, E., Mittal, M. & Sau, A. K. Understanding the lower GMP formation in large GTPase hGBP-2 and role of its individual domains in regulation of GTP hydrolysis. The FEBS journal 286, 4103–4121, doi:10.1111/febs.14957 (2019).

30 Wittinghofer, A. & Vetter, I. R. Structure-function relationships of the G domain, a canonical switch motif. Annual review of biochemistry 80, 943–971 (2011).

31 Walker, J. E., Saraste, M., Runswick, M. J. & Gay, N. J. Distantly related sequences in the alpha- and beta-subunits of ATP synthase, myosin, kinases and other ATP-requiring enzymes and a common nucleotide binding fold. The EMBO journal 1, 945–951, doi:10.1002/j.1460-2075.1982.tb01276.x (1982).

32 Scheffzek, K. et al. The Ras-RasGAP complex: structural basis for GTPase activation and its loss in oncogenic Ras mutants. Science 277, 333–338, doi:10.1126/science.277.5324.333 (1997).

33 Praefcke, G. J. et al. Identification of residues in the human guanylate-binding protein 1 critical for nucleotide binding and cooperative GTP hydrolysis. Journal of molecular biology 344, 257–269, doi:10.1016/j.jmb.2004.09.026 (2004).

34 Otwinowski, Z. M., W. Processing of X-ray diffraction data collected in oscillation mode. Methods in Enzymology 276, 307–326, doi:http://dx.doi.org/10.1016/S0076-6879(97)76066-X (1997).

35 Adams, P. D. et al. PHENIX: a comprehensive Python-based system for macromolecular structure solution. Acta Crystallogr D Biol Crystallogr 66, 213–221, doi:10.1107/S0907444909052925 (2010).

36 Emsley, P. & Cowtan, K. Coot: model-building tools for molecular graphics. Acta Crystallogr D Biol Crystallogr 60, 2126–2132, doi:10.1107/S0907444904019158 (2004).

