## Supplementary Figures for "Crystal structures reveal nucleotide-induced conformational changes in G motifs and distal regions in guanylate-binding protein 2"

```
β-1 β0 α0 β1 α1
sp|P32456|GBP2_HUMAN    ---------------MAPEINLPGPMSLIDNTKGQLVVNPEALKILSAITQPVVVVAIVGLYRTGKSYLMNKLAG   60
sp|P32455|GBP1_HUMAN    ---------------MASEIHMTGPMCLIENTNGRLMANPEALKILSAITQPMVVVAIVGLYRTGKSYLMNKLAG   60
sp|Q9H0R5|GBP3_HUMAN    ---------------MAPEIHMTGPMCLIENTNGELVANPEALKILSAITQPVVVVAIVGLYRTGKSYLMNKLAG   60
sp|Q96PP9|GBP4_HUMAN    MGERTLHAAVPTPGYPESESIMMAPICLVENQEEQLTVNSKALEILDKISQPVVVVAIVGLYRTGKSYLMNRLAG   75
sp|Q96PP8|GBP5_HUMAN    ---------------MALEIHMSDPMCLIENFNEQLKVNQEALEILSAITQPVVVVAIVGLYRTGKSYLMNKLAG   60
sp|Q6ZN66|GBP6_HUMAN    ---------------MESGPKMLAPVCLVENNNEQLLVNQQAIQILEKISQPVVVVAIVGLYRTGKSYLMNHLAG   60
sp|Q8N8V2|GBP7_HUMAN    ---------------MASEIHMPGPVCLTENTKGHLVVNSEALEILSAITQPVVVVAIVGLYRTGKSYLMNKLAG   60
                                               :  *:.* :* : .* .* :*::**. *:**:*****************:***

β2 β3 α2 β4
sp|P32456|GBP2_HUMAN    KKNGFSLGSTVKSHTKGIWMWCVPHPKKPEHTLVLLDTEGLGDIEKGDNENDSWIFALAILLSSTFVYNSMGTIN   135
sp|P32455|GBP1_HUMAN    KKKGFSLGSTVQSHTKGIWMWCVPHPKKPGHILVLLDTEGLGDVEKGDNQNDSWIFALAVLLSSTFVYNSIGTIN   135
sp|Q9H0R5|GBP3_HUMAN    KNKGFSLGSTVKSHTKGIWMWCVPHPKKPEHTLVLLDTEGLGDVKKGDNQNDSWIFTLAVLLSSTLVYNSMGTIN   135
sp|Q96PP9|GBP4_HUMAN    KRNGFPLGSTVQSETKGIWMWCVPHLSKPNHTLVLLDTEGLGDVEKSNPKNDSWIFALAVLLSSSFVYNSVSTIN   150
sp|Q96PP8|GBP5_HUMAN    KNKGFSVASTVQSHTKGIWIWCVPHPNWPNHTLVLLDTEGLGDVEKADNKNDIQIFALALLLSSTFVYNTVNKID   135
sp|Q6ZN66|GBP6_HUMAN    QNHGFPLGSTVQSETKGIWMWCVPHPSKPNHTLVLLDTEGLGDVEKGDPKNDSWIFALAVLLCSTFVYNSMSTIN   135
sp|Q8N8V2|GBP7_HUMAN    KNKGFPLGCTVKSETKGIWMWCVPHPSKPNHTLILLDTEGLGDMEKSDPKSDSWIFALAVLLSSSFVYNSMGTIN   135
                        :.:** :..**:*.*****:***** . * * *:*********::*.: :.*  **:**:**.*::***::..*:

α3 α3' β5 α4
sp|P32456|GBP2_HUMAN    QQAMDQLHYVTELTDRIKANSSPGN--NSVDDSADFVSFFPAFVWTLRDFTLELEVDGEPITADDYLELSLKLRK   208
sp|P32455|GBP1_HUMAN    QQAMDQLYYVTELTHRIRSKSSPDENENEVEDSADFVSFFPDFVWTLRDFSLDLEADGQPLTPDEYLTYSLKLKK   210
sp|Q9H0R5|GBP3_HUMAN    QQAMDQLYYVTELTHRIRSKSSPDENEN--EDSADFVSFFPDFVWTLRDFSLDLEADGQPLTPDEYLEYSLKLTQ   208
sp|Q96PP9|GBP4_HUMAN    HQALEQLHYVTELAELIRAKSCPRP--DEAEDSSEFASFFPDFIWTVRDFTLELKLDGNPITEDEYLENALKLIP   223
sp|Q96PP8|GBP5_HUMAN    QGAIDLLHNVTELTDLLKARNSPDL--DRVEDPADSASFFPDLVWTLRDFCLGLEIDGQLVTPDEYLENSLRPKQ   208
sp|Q6ZN66|GBP6_HUMAN    HQALEQLHYVTELTELIKAKSSPRP--DGVEDSTEFVSFFPDFLWTVRDFTLELKLNGHPITEDEYLENALKLIQ   208
sp|Q8N8V2|GBP7_HUMAN    HQALEQLHYVTELTELIRAKSCPRP--DEVEDSSEFVSFFPDFIWTVRDFTLELKLDGHPITEDEYLENALKLIS   208
                        :*:: *: ****:. :::...*    :   :* :: .**** ::**:*** * *: :*. :* *:**  :*:

α4' β6 α5
sp|P32456|GBP2_HUMAN    GTDKKSKSFNDPRLCIRKFFPKRKCFVFDWPAPKK-YLAHLEQLKEEELNPDFIEQVAEFCSYILSHSNVKTLSG   282
sp|P32455|GBP1_HUMAN    GTSQKDETFNLPRLCIRKFFPKKKCFVFDRPVHRR-KLAQLEKLQDEELDPEFVQQVADFCSYIFSNSKTKTLSG   284
sp|Q9H0R5|GBP3_HUMAN    GTSQKDKNFNLPRLCIRKFFPKKKCFVFDLPIHRR-KLAQLEKLQDEELDPEFVQQVADFCSYIFSNSKTKTLSG   282
sp|Q96PP9|GBP4_HUMAN    GKNPKIQNSNMPRECIRHFFRKRKCFVFDRPTNDKQYLNHMDEVPEENLERHFLMQSDNFCSYIFTHAKTKTLRE   298
sp|Q96PP8|GBP5_HUMAN    GSDQRVQNFNLPRLCIQKFFPKKKCFIFDLPAHQK-KLAQLETLPDDELEPEFVQQVTEFCSYIFSHSMTKTLPG   282
sp|Q6ZN66|GBP6_HUMAN    GNNPRVQTSNFPRECIRRFFPKRKCFVFDRPTNDKDLLANIEKVSEKQLDPKFQEQTNIFCSYIFTHARTKTLRE   283
sp|Q8N8V2|GBP7_HUMAN    GKNPQIQNSNKPREWIRHFFPKQKCFVFDRPINDKKLLLHVEEVREDQLDSNFQMQSENFCSYIFTHAKTKTLRE   283
                        *.. : :. * **  *::** *:***:** *   :  * ::: : :.:*: .*  *   *****:::: .***

α6 α7 α8
sp|P32456|GBP2_HUMAN    GIPVNGPRLESLVLTYVNAISSGDLPCMENAVLALAQIENSAAVEKAIAHYEQQMGQKVQLPTETLQELLDLHRD   357
sp|P32455|GBP1_HUMAN    GIQVNGPRLESLVLTYVNAISSGDLPCMENAVLALAQIENSAAVQKAIAHYEQQMGQKVQLPTETLQELLDLHRD   359
sp|Q9H0R5|GBP3_HUMAN    GIKVNGPRLESLVLTYINAISRGDLPCMENAVLALAQIENSAAVQKAIAHYDQQMGQKVQLPAETLQELLDLHRV   357
sp|Q96PP9|GBP4_HUMAN    GIIVTGKRLGTLVVTYVDAINSGAVPCLENAVTALAQLENPAAVQRAADHYSQQMAQQLRLPTDTLQELLDVHAA   373
sp|Q96PP8|GBP5_HUMAN    GIMVNGSRLKNLVLTYVNAISSGDLPCIENAVLALAQRENSAAVQKAIAHYDQQMGQKVQLPMETLQELLDLHRT   357
sp|Q6ZN66|GBP6_HUMAN    GITVTGNRLGTLAVTYVEAINSGAVPCLENAVITLAQRENSAAVQRAADYYSQQMAQRVKLPTDTLQELLDMHAA   358
sp|Q8N8V2|GBP7_HUMAN    GILVTGNRLGMLVETYLDAINSGATPCLENAMAVLAQCENSAAVQRAANHYSQQMAQQVRFPTDTLQELLDVHAV   358
                        ** *.* **  *. **::**. *  **:***: .*** ** ***::*  :*.***.*::::* :*******:*

α8 α9
sp|P32456|GBP2_HUMAN    SEREAIEVFMKNSFKDVDQMFQRKLGAQLEARRDDFCKQNSKASSDCCMALLQDIFGPLEEDVKQGTFSKPGGYR   432
sp|P32455|GBP1_HUMAN    SEREAIEVFIRSSFKDVDHLFQKELAAQLEKKRDDFCKQNQEASSDRCSALLQVIFSPLEEEVKAGIYSKPGGYR   434
sp|Q9H0R5|GBP3_HUMAN    SEREATEVYMKNSFKDVDHLFQKKLAAQLDKKRDDFCKQNQEASSDRCSALLQVIFSPLEEEVKAGIYSKPGGYC   432
sp|Q96PP9|GBP4_HUMAN    CEREAIAVFMEHSFKDENHEFQKKLVDTIEKKKGDFVLQNEEASAKYCQAELKRLSEHLTESILRGIFSVPGGHN   438
sp|Q96PP8|GBP5_HUMAN    SEREAIEVFMKNSFKDVDQSFQKELETLLDAKQNDICKRNLEASSDYCSALLKDIFGPLEEAVKQGIYSKPGGHN   432
sp|Q6ZN66|GBP6_HUMAN    CEREAIAIFMEHSFKDENQEFQKKFMETTMNKKGDFLLQNEESSVQYCQAKLNELSKGLMESISAGSFSVPGGHK   433
sp|Q8N8V2|GBP7_HUMAN    CEREAIAVFMEYSFKDKSQEFQKKLVDTMEKKKEDFVLQNEEASAKYCQAELKRLSELLTESISRGTFFVPGGHN   433
                        .****  :::. **** .: **:::      :: *:  :* ::* . * * *: :   * * :  * :  ***:

α10 α11 α12
sp|P32456|GBP2_HUMAN    LFTQKLQELKNKYYQVPRKGIQAKEVLKKYLESKEDVADALLQTDQSLSEKEKAIEVERIKAESAEAAKKMLEEI   507
sp|P32455|GBP1_HUMAN    LFVQKLQDLKKKYYEEPRKGIQAEEILQTYLKSKESMTDAILQTDQTLTEKEKEIEVERVKAESAQASAKMLQEM   509
sp|Q9H0R5|GBP3_HUMAN    LFIQKLQDLEKKYYEEPRKGIQAEEILQTYLKSKESVTDAILQTDQILTEKEKEIEVECVKAESAQASAKMVEEM   507
sp|Q96PP9|GBP4_HUMAN    LYLEEKKQVEWDYKLVPRKGVKANEVLQNFLQSQVVVEESILQSDKALTAGEKAIAAERAMKEAAEKEQELLREK   523
sp|Q96PP8|GBP5_HUMAN    LFIQKTEELKAKYYREPRKGIQAEEVLQKYLKSKESVSHAILQTDQALTETEKKKEAQVKAEAEKAEAQRLAAI   507
sp|Q6ZN66|GBP6_HUMAN    LYMETKERIEQDYWQVPRKGVKAKEVFQRFLESQMVIEESILQSDKALTDREKAVAVDRAKKEAAEKEQELLKQK   508
sp|Q8N8V2|GBP7_HUMAN    IYLEAKKKIEQDYTLVPRKGVKADEVLQSFLQSQVVIEESILQSDKALTAGEKAIAAKQAKKEAAEKEQELLRQK   508
                        :: :  : :: .*    ****::*.*::: :*:*:  : .::**:*: *:  **          *: :   : :

α12 α13
sp|P32456|GBP2_HUMAN    QKKNEEMMEQKEKSYQEHVKQLTEKMERDRAQLMAEQEKTLALKLQEQERLLKEGFENESKRLQKDIWDIQMRSK   582
sp|P32455|GBP1_HUMAN    QRKNEQMMEQKERSYQEHLKQLTEKMENDRVQLLKEQERTLALKLQEQEQLLKEGFQKESRIMKNEIQDLQTKMR   584
sp|Q9H0R5|GBP3_HUMAN    QIKYQQMMEEKEKSYQEHVKQLTEKMERERAQLLEEQEKTLTSKLQEQARVLKERCQGESTQLQNEIQKLQKTLK   582
sp|Q96PP9|GBP4_HUMAN    QKEQQQMMEAQERSFQEYMAQMEKKLEEERENLLREHERLLKHKLKVQEEMLKEEFQKKSEQLNKEINQLKEKIE   598
sp|Q96PP8|GBP5_HUMAN    QRQNEQMMQERERLHQEQVRQME----IAKQNWLAEQQKMQEQQMQEQAAQLSTTFQAQNRSLLSELQHAQRTVN   578
sp|Q6ZN66|GBP6_HUMAN    LQEQQQQMEAQDKSRKENIAQLKEKLQMEREHLLREQIMMLEHTQKVQNDWLHEGFKKKYEEMNAEISQFKRMID   583
sp|Q8N8V2|GBP7_HUMAN    QKEQQQMMEAQERSFQENIAQLKKKMERERENYMRELRKMLSHKMKVLEELLTEGFKEIFESLNEEINRLKEQIE   583
                          : :: *: :::  :* : *:        : : : *          :     *    :     :  ::   :

sp|P32456|GBP2_HUMAN    SLEP--ICNIL------------------------------------------ 591
sp|P32455|GBP1_HUMAN    RRK---ACTIS------------------------------------------ 592
sp|Q9H0R5|GBP3_HUMAN    KKT---KRYMSHKLKI------------------------------------- 595
sp|Q96PP9|GBP4_HUMAN    STKNEQLR-LLKILDMASNIMIVTLPGASKLLGVGTKYLGSRI---------- 640
sp|Q96PP8|GBP5_HUMAN    NDDP--CVLL------------------------------------------- 586
sp|Q6ZN66|GBP6_HUMAN    TTKNDDTPWIARTLDNLADELTAILSAPAKLIGHGVKGVSSLFKKHKLPF----- 633
sp|Q8N8V2|GBP7_HUMAN    AAENEEPSVFSQILDVAGSIFIAALPGAAKLVDLGMKILSSLCNRLRNPGKKIIS 638
                               :
```

G1/P loop

G2/Switch I

G3/Switch II

G4

**Figure S1. Sequence alignment of different guanylate binding proteins in human showing conserved G1 (lavender), G2 (aqua), G3 (orange) and G4 (dark green) motifs.** The secondary structural elements are represented at the top of the sequence, arrows represent beta sheets, rectangular boxes represent alpha helices, and the straight lines represent loops.

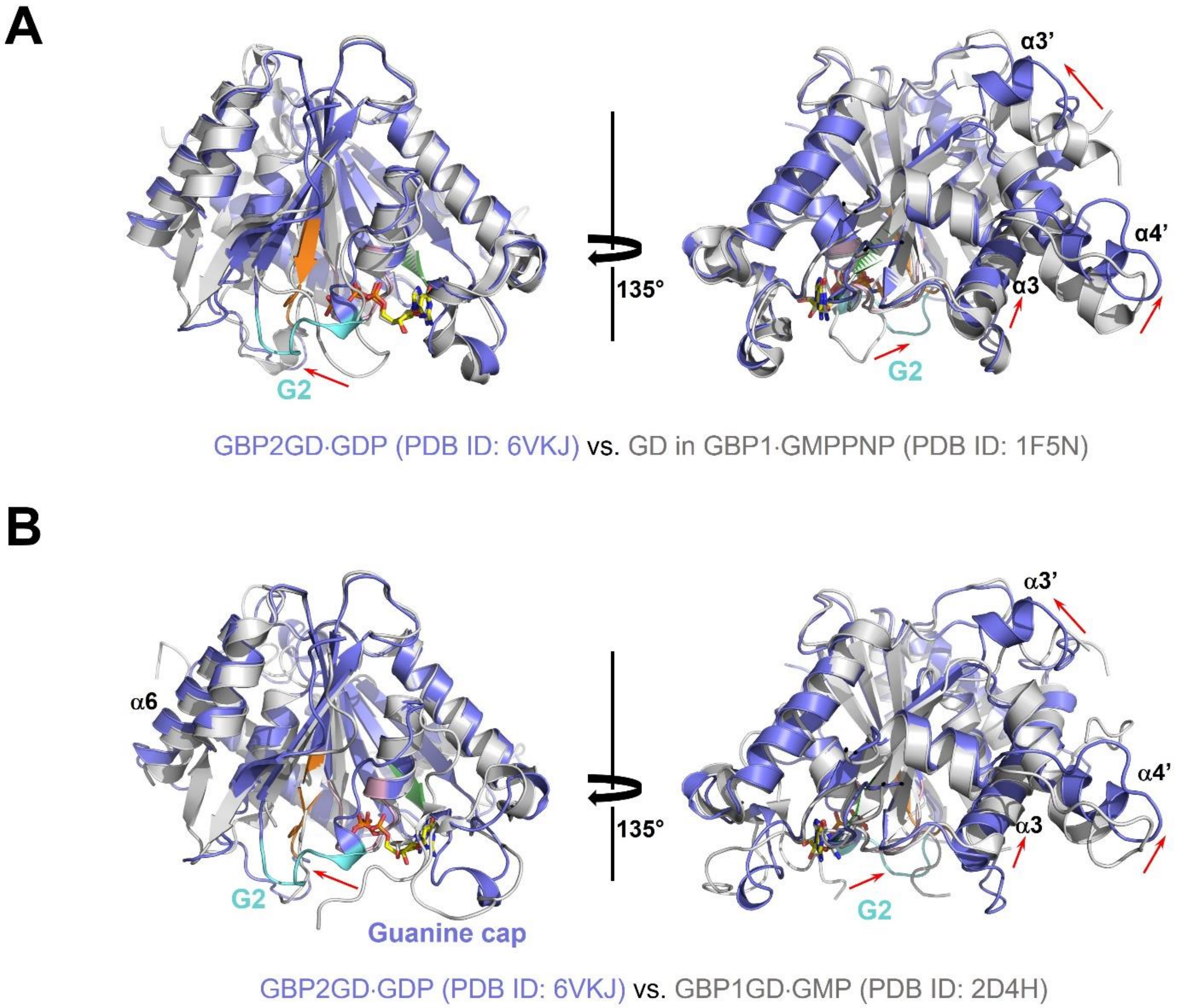


**Figure S2. GBP2GD·GDP displays substantial conformational changes from GBP1GD (in FL)·GMPPNP and GBP1GD·GMP.** (**A**) Superposition of GBP2GD·GDP and GBP1GD·GMPPNP structures in two views. GBP2GD is colored as in **Fig. 2A** while GBP1 is colored in gray. Red arrows point to the regions with substantial conformational changes. (**B**) Superposition of GBP2GD·GDP and GBP1GD·GMP structures in two views. GBP2GD is colored as in **Fig. 2A** while GBP1 is colored in gray. Red arrows point to the regions with substantial conformational changes.

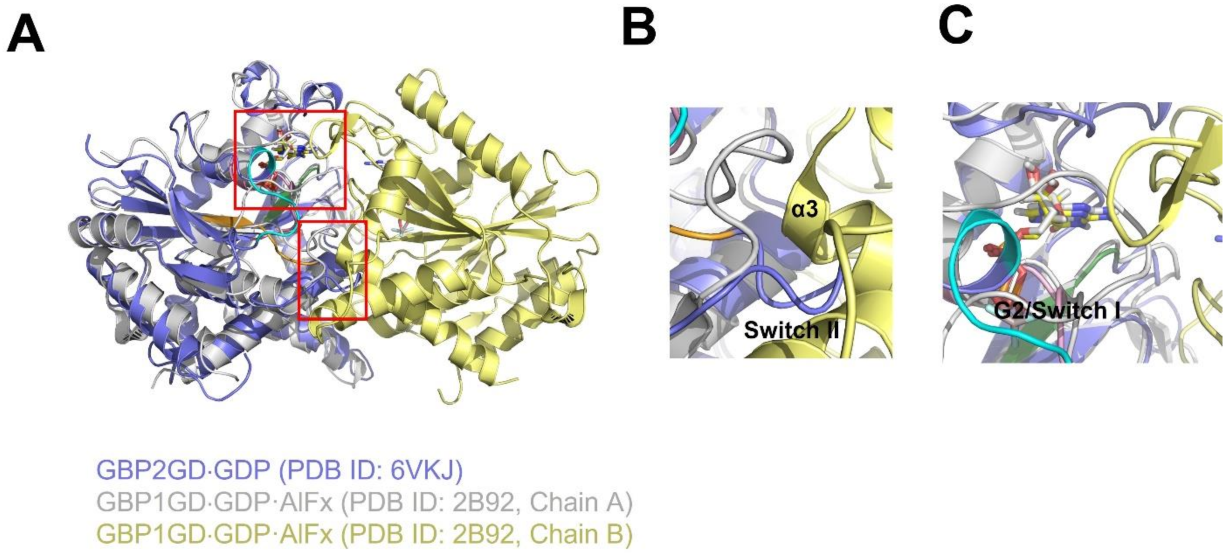


**Figure S3. GBP2GD·GDP is incompatible with dimerization.** (**A**) Superposition of GBP2GD·GDP structure with dimeric GBP1GD structure reveal clashes. The two GBP1GD molecules are colored in gray and pale yellow. The regions that cause clash or reduce interaction are boxed in red. (**B**) GDP binding causes switch II region to clash with α3 of the other protomer. (**C**) Retraction of switch I region from dimer interface in GDP-bound form reduces interaction with the other protomer.

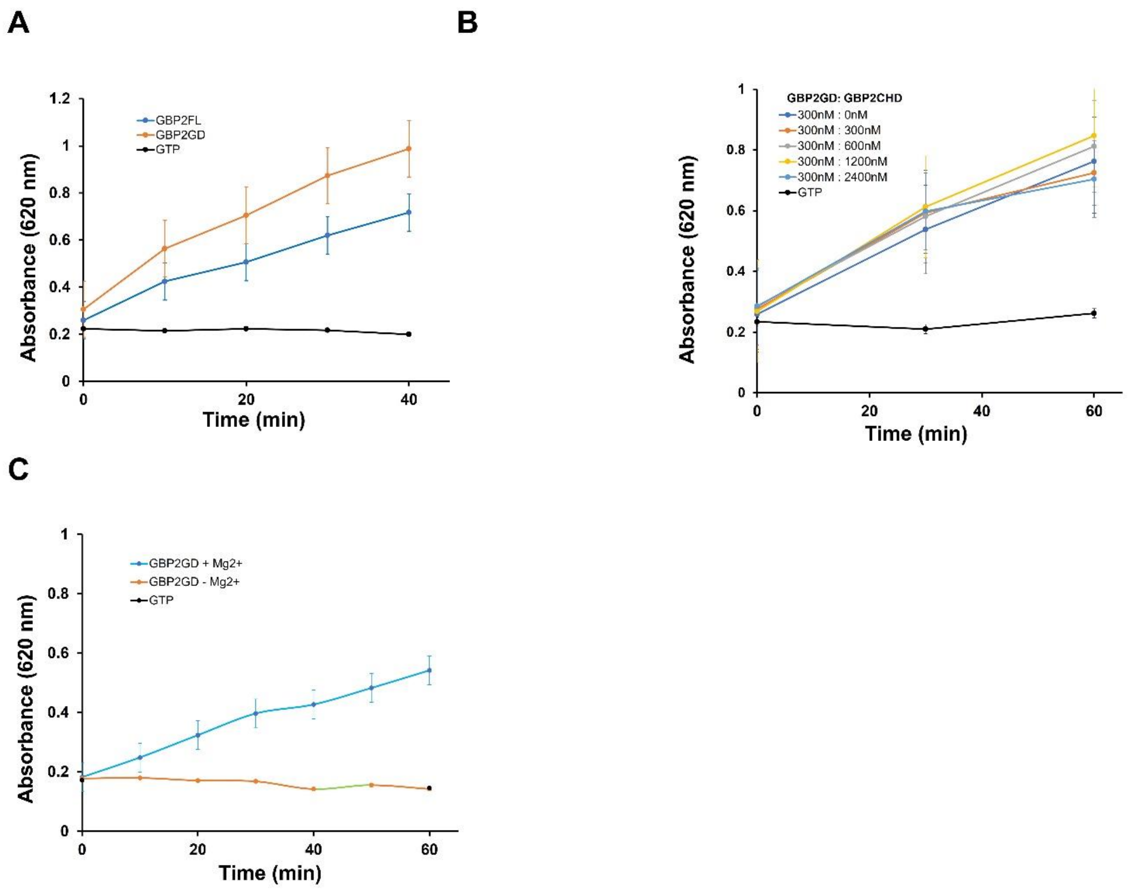


**Figure S4** (**A**) Comparison of GTPase activities of full-length GBP2 (GBP2FL) and GBP2GD. (**B**) Measurement of GBP2GD activity in increasing amount of GBP2CHD. (**C**) Dependence of GBP2GD activity on magnesium.

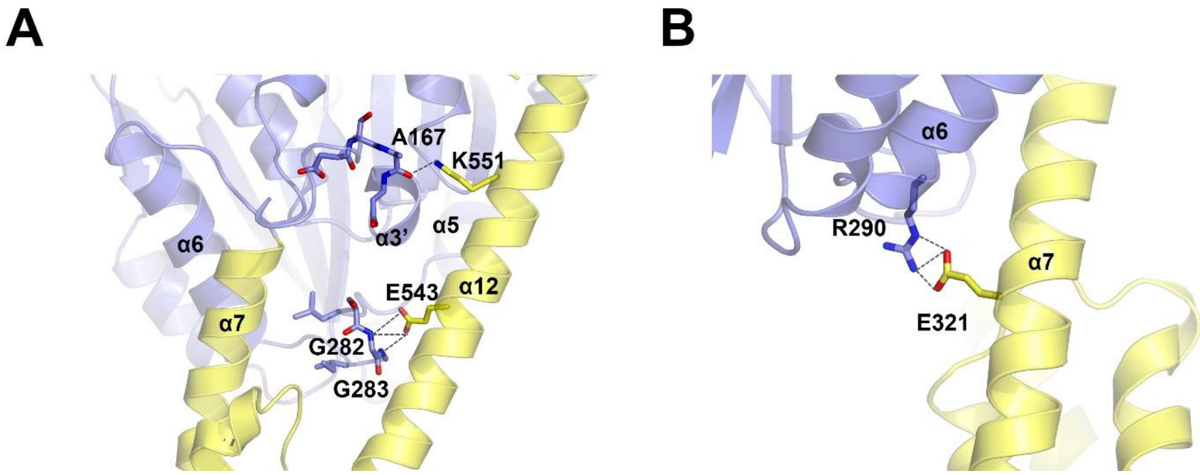


**Figure S5. Additional interactions at the GD:CHD interface.** (**A**) Top: K551 on α12 interacting with the carbonyl oxygen of A167; Bottom: E543 interacting with the mainchain amine groups of G282 and G283. α13 is removed for clarity reasons. (B) The interaction between R290 on α6 and E321 on α7. GBP2 is colored as in Figure 4 with GD colored in blue and CHD in yellow. Dashed lines represent hydrogen bonds or salt bridges.

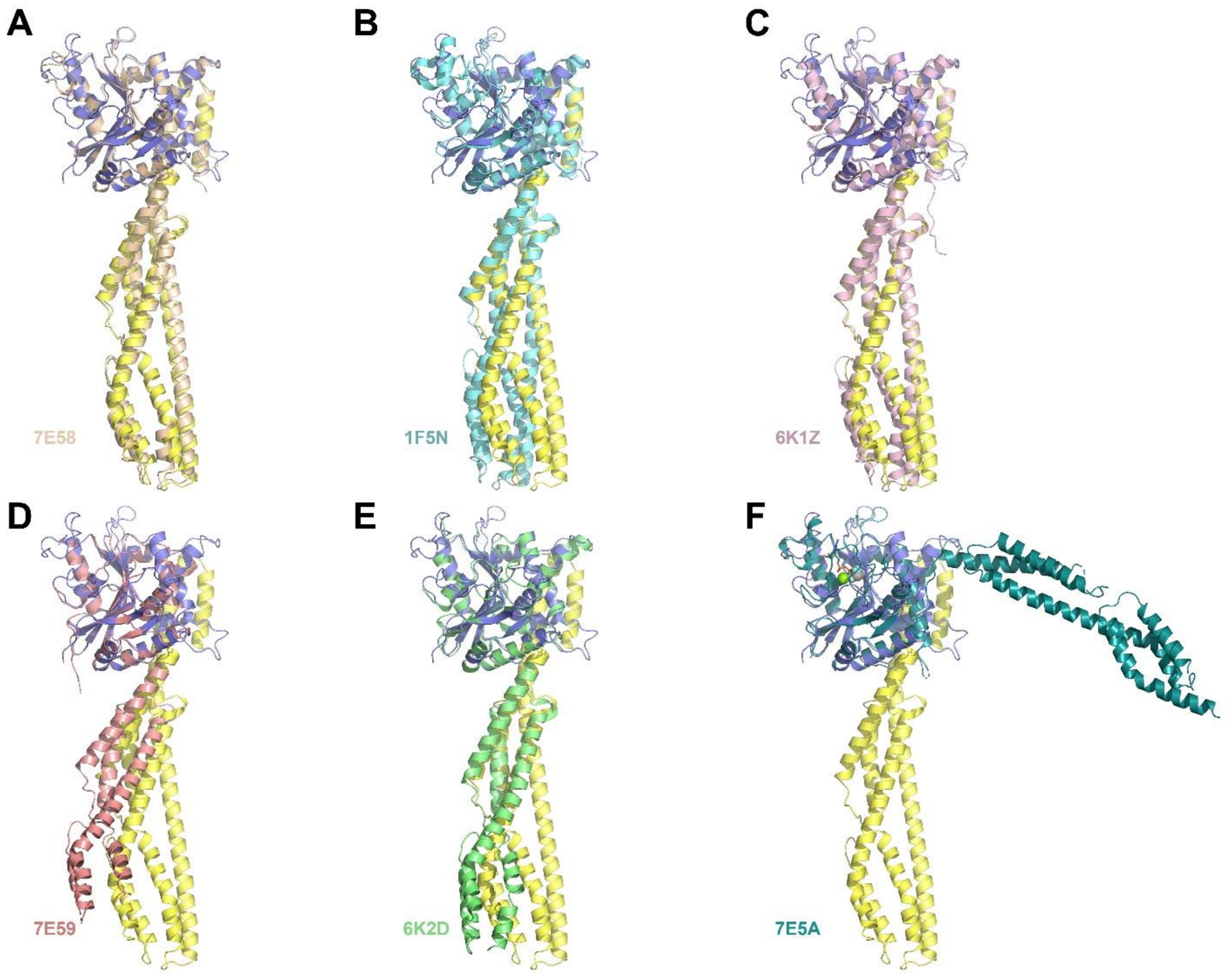


**Figure S6. Comparison of the CHD conformations in GBP structures.** GBP2$^{K51A}$ structure (blue and yellow) superposed with full-length GBP proteins (A-C) or truncated GBP proteins (D-F). Only the GD domains are superposed to accentuate the swinging movement of the CHD. **(A)** GBP2$^{K51A}$ superposed with nucleotide-free full-length GBP2 structure (pale orange, PDB ID 7E58). (**B**) GBP2$^{K51A}$ superposed with GMPPNP-bound full-length GBP1 structure (cyan, PDB ID 1F5N). (**C**) GBP2$^{K51A}$ superposed with nucleotide-free farnesylated full-length GBP1 structure (pink, PDB ID 6K1Z). (**D**) GBP2$^{K51A}$ superposed with nucleotide-free GBP1Δα12-α13 structure (in complex with IpaH9.8) (lime green, PDB ID 6K2D). (**E**) GBP2$^{K51A}$ superposed with GBP5Δα12-α13 structure (salmon, PDB ID 7E59). (**F**) GBP2$^{K51A}$ superposed with GBP5Δα12-α13 structure in the dimeric conformation (teal, PDB ID 7E5A).

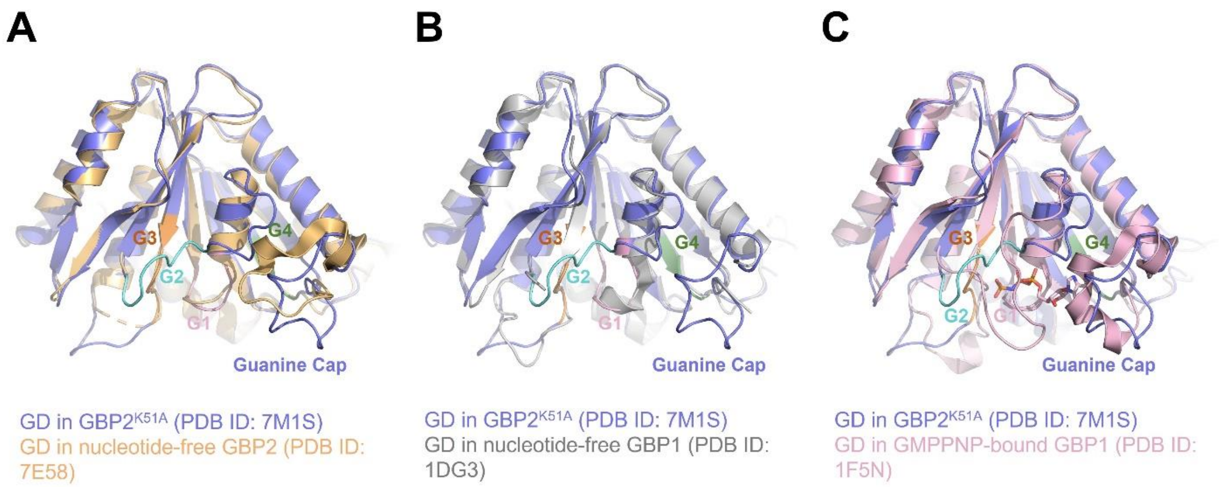


**Figure S7. Comparison of G domain conformations in nucleotide free full-length GBP1 and GBP2 structures. (A)** GD conformation in GBP2$^{K51A}$ structure (blue) superposed with GD domain in GBP2 structure (light orange, PDB ID: 7E58). (**B**) GD conformation in GBP2$^{K51A}$ structure (blue) superposed with GD domain in GBP1 structure (gray, PDB ID: 1DG3). (**C**) GD conformation in GBP2$^{K51A}$ structure (blue) superposed with GD domain in GMPPNP-bound GBP1 structure (pink, PDB ID: 1F5N). The GMPPNP molecule is shown as stick model. In the GBP2$^{K51A}$ structure, the G1-G4 motifs are colored in pink, cyan, orange, and deep green as in Figure 2.

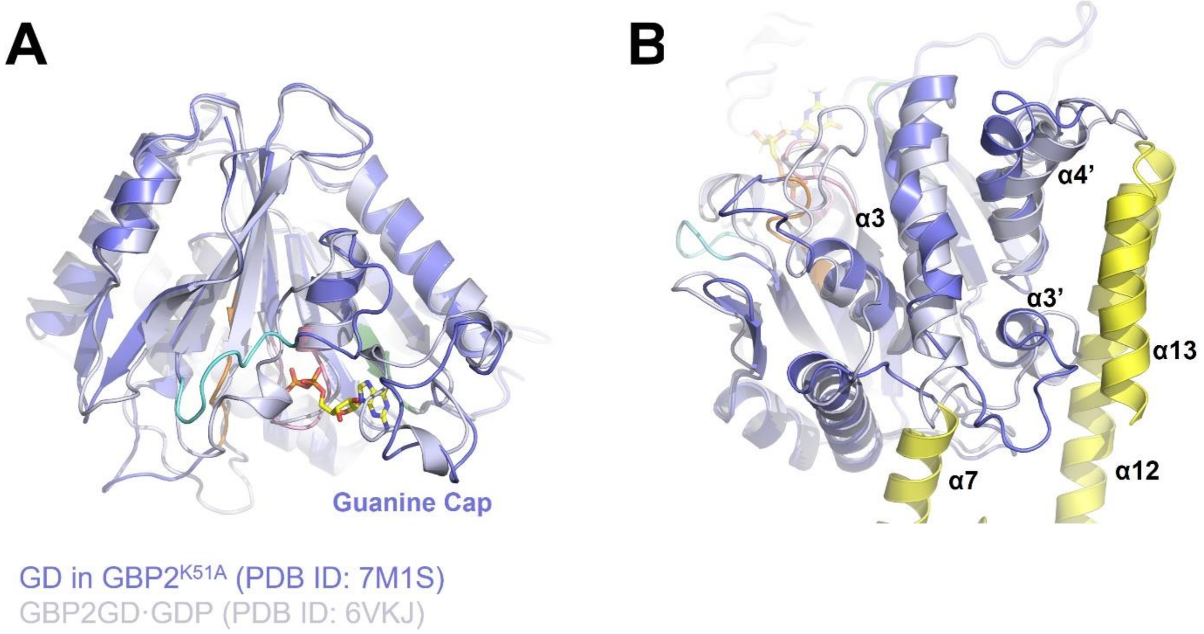


**Figure S8. Comparison of G domain conformations full-length GBP1$^{K51A}$ and GBP2GD with GDP structures. (A)** GD conformation in GBP2$^{K51A}$ structure (blue) superposed with GD domain in GBP2GD·GDP structure (light blue, PDB ID: 6VKJ). (B) Conformational differences in the CHD-contacting region between GBP2$^{K51A}$ structure and GBP2GD·GDP structure. In the GBP1$^{K51A}$ structure, the G1-G4 motifs are colored in pink, cyan, orange, and deep green as in Figure 2. GDP is shown as sticks.
