## Supplementary Table 1 for "Crystal structures reveal nucleotide-induced conformational changes in G motifs and distal regions in guanylate-binding protein 2"

|  | **GBP2GD·GDP** | **GBP2FL (K51A)** |
| --- | --- | --- |
| **Data Collection** |  |  |
| Wavelength (Å) | 0.920126 | 0.920126 |
| Resolution range (Å) | 45.17 - 2.105 (2.181 - 2.105) | 45.27 - 2.91 (3.02 - 2.91) |
| Space group | P 41 21 2 | C 2 2 21 |
| Unit cell (a, b, c, Å)  (α, β, γ, °) | 87.951 87.951 105.279  90 90 90 | 59.631 140.569 240.079  90 90 90 |
| Unique reflections | 24391 (2231) | 22422 (1997) |
| Multiplicity | 16.7 (10.4) | 9.4 (7.5) |
| Completeness (%) | 99.31 (93.11) | 98.81 (89.49) |
| Mean I/sigma(I) | 24.61 (1.42) | 12.16 (1.21) |
| Wilson B-factor | 39.80 | 84.99 |
| R-merge | 0.244 (1.334) | 0.173 (1.594) |
| R-meas | 0.251 (1.404) | 0.182 (1.694) |
| R-pim | 0.059 (0.432) | 0.056 (0.559) |
| CC1/2 | 0.993 (0.697) | 0.998 (0.747) |
| CC* | 0.998 (0.906) | 0.999 (0.911) |
| **Refinement** |  |  |
| Reflections used in refinement | 24389 (2231) | 22383 (1984) |
| Reflections used for R-free | 2000 (183) | 1995 (178) |
| R-work | 0.2275 (0.3213) | 0.2451 (0.4230) |
| R-free | 0.2390 (0.3348) | 0.2736 (0.4278) |
| Number of non-hydrogen atoms | 2552 | 4729 |
| macromolecules | 2394 | 4608 |
| ligands | 28 | 5 |
| solvent | 130 | 116 |
| Protein residues | 305 | 577 |
| RMS (bonds, Å) | 0.017 | 0.011 |
| RMS (angles, °) | 1.31 | 1.54 |
| Ramachandran favored (%) | 97.03 | 93.91 |
| Ramachandran allowed (%) | 2.97 | 5.91 |
| Ramachandran outliers (%) | 0.00 | 0.17 |
| Rotamer outliers (%) | 1.85 | 1.17 |
| Clashscore | 18.81 | 15.58 |
| Average B-factor | 54.09 | 100.55 |
| macromolecules | 55.49 | 99.89 |
| ligands | 46.08 | 112.01 |
| solvent | 30.00 | 126.24 |
| **Number of TLS groups** | 1 | 1 |
