## Supplementary Table 2 for "Crystal structures reveal nucleotide-induced conformational changes in G motifs and distal regions in guanylate-binding protein 2"

| **Constructs used** | **Primer sequence** |
| --- | --- |
| GBP2FL | Forward: CGC**GGATCC**GCTCCAGAGATCAACTTGCCG  Reverse: ACGC**GTCGAC**TTAGAGTATGTTACATATTGGCTCC |
| GBP2GD | Forward: CGC**GGATCC**GCTCCAGAGATCAACTTGCCG  Reverse: ACGC**GTCGAC**TTAGCAGGGTAGATCCCCACTG |
| GBP2CHD | Forward: CGC**GGATCC**GAGAACGCAGTCCTGGCCTTG  Reverse: ACGC**GTCGAC**TTAGAGTATGTTACATATTGGCTCC |
| GBP2FL^K51A^ | Forward: GGCCTCTATCGCACAGGCGCATCCTACCTGATGAACAA  Reverse: TTGTTCATCAGGTAGGATGCGCCTGTGCGATAGAGGCC |
| GBP2FL^R48A^ | Forward: CGATTGTGGGCCTCTATGCGACAGGCAAATCCTACCTG  Reverse: CAGGTAGGATTTGCCTGTCGCATAGAGGCCCACAATCG |
| GBP2FL^S73A^ | Forward: CTCTAGGCTCCACAGTGAAGGCGCACACCAAGGGAATCTGG  Reverse: CCAGATTCCCTTGGTGTGCGCCTTCACTGTGGAGCCTAGAG |
| GBP2FL^E99Q^ | Forward: CCCTAGTTCTGCTCGACACTCAGGGCCTGGGAGATATAGAGAAG  Reverse: CTTCTCTATATCTCCCAGGCCGTCAGTGTCGAGCAGAACTAGGG |
| GBP2FL^E99D^ | Forward: CCCTAGTTCTGCTCGACACTGATGGCCTGGGAGATATAGAGAAG  Reverse: CTTCTCTATATCTCCCAGGCCCTAAGTGTCGAGCAGAACTAGGG |
| GBP2FL^E99L^ | Forward: CCCTAGTTCTGCTCGACACTCTGGGCCTGGGAGATATAGAGAAG  Reverse: CTTCTCTATATCTCCCAGGCCGACAGTGTCGAGCAGAACTAGGG |
